# First evidence of production of the lantibiotic nisin P

**DOI:** 10.1101/827204

**Authors:** Enriqueta Garcia-Gutierrez, Paula M. O’Connor, Gerhard Saalbach, Calum J. Walsh, James W. Hegarty, Caitriona M. Guinane, Melinda J. Mayer, Arjan Narbad, Paul D. Cotter

**Author notes:** **Corresponding author:** Professor Arjan Narbad.

## Abstract

Nisin P is a natural nisin variant, the genetic determinants for which were previously identified in the genomes of two *Streptococcus* species, albeit with no confirmed evidence of production. Here we describe *Streptococcus agalactiae* DPC7040, a human fecal isolate, which exhibits antimicrobial activity against a panel of gut and food isolates by virtue of producing nisin P. Nisin P was purified, and its predicted structure was confirmed by nanoLC-MS/MS, with both the fully modified peptide and a variant without rings B and E being identified. Additionally, we compared its spectrum of inhibition and minimum inhibitory concentration (MIC) with that of nisin A and its antimicrobial effect in a fecal fermentation in comparison with nisin A and H. We found that its antimicrobial activity was less potent than nisin A and H, and we propose a link between this reduced activity and the peptide structure.

## Introduction

Nisin is a small peptide with antimicrobial activity against a wide range of pathogenic bacteria. It was originally sourced from a *Lactococcus lactis* subsp. *lactis* isolated from a dairy product ^1^ and is classified as a class I bacteriocin, as it is ribosomally synthesized and post-translationally modified ^2^. Nisin has been studied extensively and has a wide range of applications in the food industry, biomedicine, veterinary and research fields ^3–6^. It is approved as a food preservative by the Food and Agriculture Organization and is Generally Recognized As Safe (GRAS) by the United States Food and Drug Administration ^4^.

Structurally, nisin is classified as a lantibiotic because it contains lanthionine (Lan), an unusual amino acid formed by two alanine residues linked by a sulphur atom through their β-carbon ^4^. Other unusual amino acids present in nisin are dehydroalanine (Dha), dehydrobutyrine (Dhb) and β-methyl-lanthionine ^4^. Nisin activity and stability are closely related to its structure and can be altered by pH changes. Increasing pH results in decreasing activity as a consequence of alterations in structure, therefore, nisin is more stable at lower pH. Nisin is heat-stable and also exhibits high stability at low temperatures, which makes it suitable for freeze-storage ^7^.

Nisin has nine reported natural variants (Figure 1, Table 1). Nisin A is the most studied nisin as it was the first one purified ^8^. Nisin Z is considered the first natural variant of nisin A; it differs in the presence of an asparagine amino acid in position 27 instead of a histidine residue ^9^. This substitution has very little effect on antimicrobial activity, thermal and pH stability compared to nisin A, but affects the solubility of the molecule, with nisin Z being more soluble at neutral pH ^10^. A study of its distribution also revealed that nisin Z is more widespread than nisin A ^10^. Nisin F, produced by *L. lactis* isolated from a fish gut, also has asparagine and valine in positions 21 and 30 ^11^. Nisin Q was identified in an environmental *L. lactis* isolate and differs from nisin A due to the presence of valine, leucine, asparagine and valine in positions 15, 21, 27 and 30, respectively ^12^. Nisin U and U2 were the first variants found to be produced by a species other than *L. lactis* ^13^; they are produced by *Streptococcus uberis* strain 42, isolated from bovine mammary secretions ^14^, and differ to nisin A by nine and ten amino acids, respectively. Nisin H, produced by *Streptococcus hyointestinalis* was the first variant isolated from a mammalian gastrointestinal (GI) tract (porcine) ^15^ and has five different amino acids at positions 1, 6, 18, 21 and 31. In 2017, a nisin O operon has been reported in *Blautia obeum* A2-162 isolated from a human GI tract ^16^. Unusually this operon encodes 4 peptides, the first 3 being identical and exhibiting similarity to nisin U, and the fourth showing the highest divergence from nisin A. Most interestingly, a further example of a multi-peptide operon from *Blautia producta* has recently been described and shown to be important in resistance against vancomycin-resistant *Enterococcus* in an animal model ^17^. The nisin P operon was previously identified in the genomes of two *Streptococcus* species ^15^ but its activity has not been reported before. Nisin P is predicted to be three amino acids shorter than nisin A at the C-terminus, making it the shortest natural nisin variant reported to date, and differs to nisin A in ten amino acids (Figure 1).

**Table 1.** Summary of nisin variants reported in the literature

| Nisin variant | First reported in | Reference |
| --- | --- | --- |
| A | *L. lactis* subsp. *lactis* | <sup>1</sup> |
| Z | *L. lactis* strain NIZO 22186 (dairy product) | <sup>9</sup> |
| Q | *L. lactis* 61-14 (Japanese river water) | <sup>12</sup> |
| F | *L. lactis* F10 (freshwater catfish) | <sup>11</sup> |
| U and U2 | *S. uberis* strain 42 (bovine mammary secretions) | <sup>13</sup> |
| P | *Streptococcus gallolyticus* subsp. *pasteurianus* | <sup>17</sup> |
| H | *S. hyointestinalis* DPC 6484 (porcine intestine) | <sup>15</sup> |
| O | *Blautia obeum* A2-162 (human GI tract) | <sup>16</sup> |

**Figure 1.**
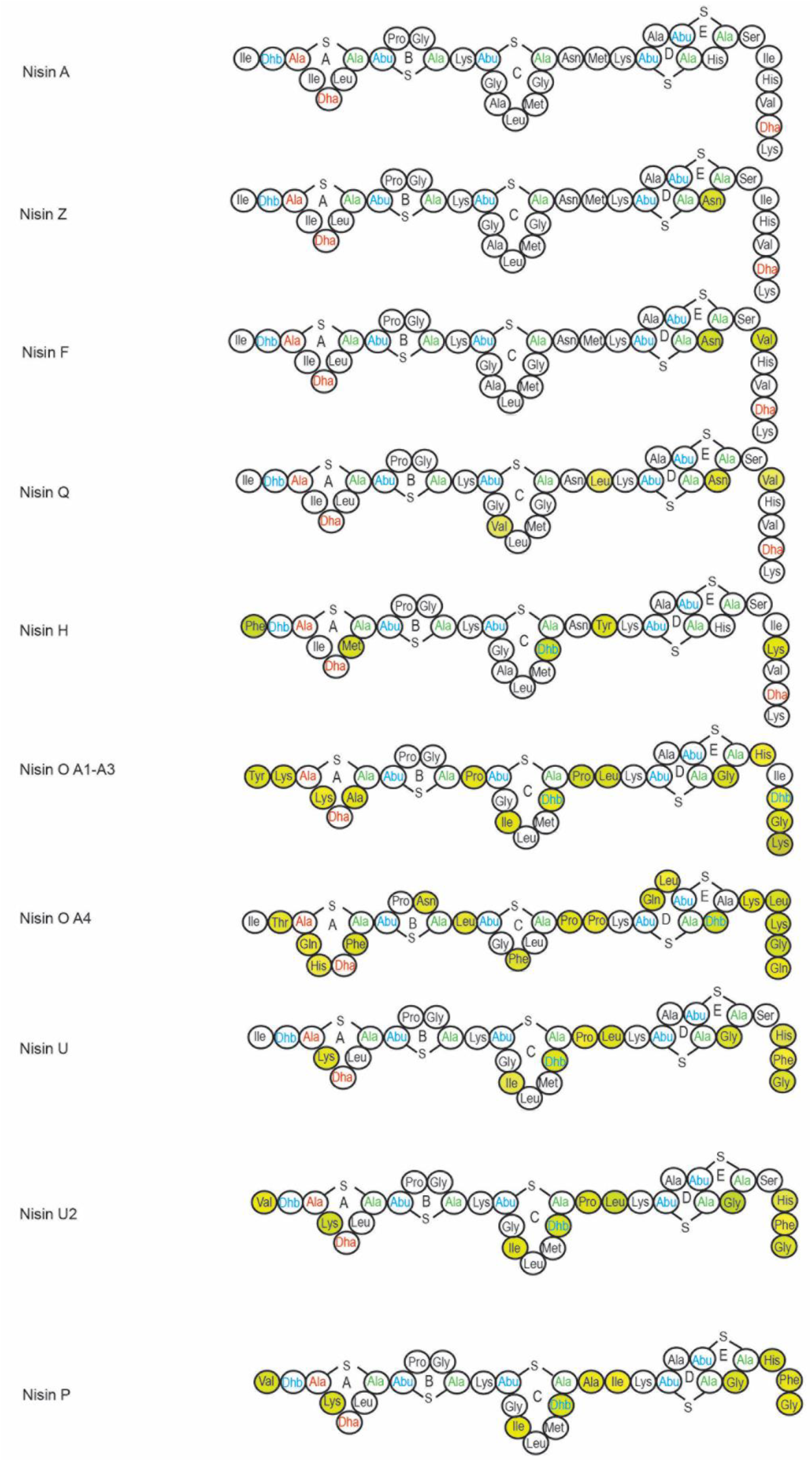
Predicted ring structures of nisin natural variants. Residues highlighted in yellow depict amino acid changes when compared to nisin A.

Here we establish that *Streptococcus agalactiae* DPC7040, isolated from human feces, contains a nisin P gene cluster and represents the first example of a strain that produces nisin P. In this study we assess the cross immunity of other nisin producers to nisin P, determine its activity *in vitro* compared to nisin A and H and investigate its ability to induce the nisin A promoter. In addition, a fecal fermentation experiment was conducted to determine how a human fecal microbiota was affected by the presence of nisin A, H and P.

## Results

### S. agalactiae *DPC7040 localization within the* S. agalactiae *species*

The constructed *S. agalactiae* phylogenetic tree showed no specificity for the body isolation site. The 13 genomes of *S. agalactiae* that were reported to have stool origin were spread over the whole species and showed no specific clustering (Figure 2A). Scoary analysis showed that there were no genes associated to the gut isolates with 100% sensitivity and specificity. However, there were genes that were significantly associated (using a Bonferroni-adjusted p value < 0.05) with stool isolates, either by presence or absence (Table S1). The PCoA constructed from accessory gene content showed some level of differential clustering between *S. agalactiae* isolates of human origin and isolates from non-human animals (Figure 2B). No annotation for LanB or LanC proteins was found in the genomes of the other stool *S. agalactiae* isolates, indicating that, among genome-sequenced strains, the nisin P operon is only present in *S. agalactiae* DPC7040.

**Figure 2.**
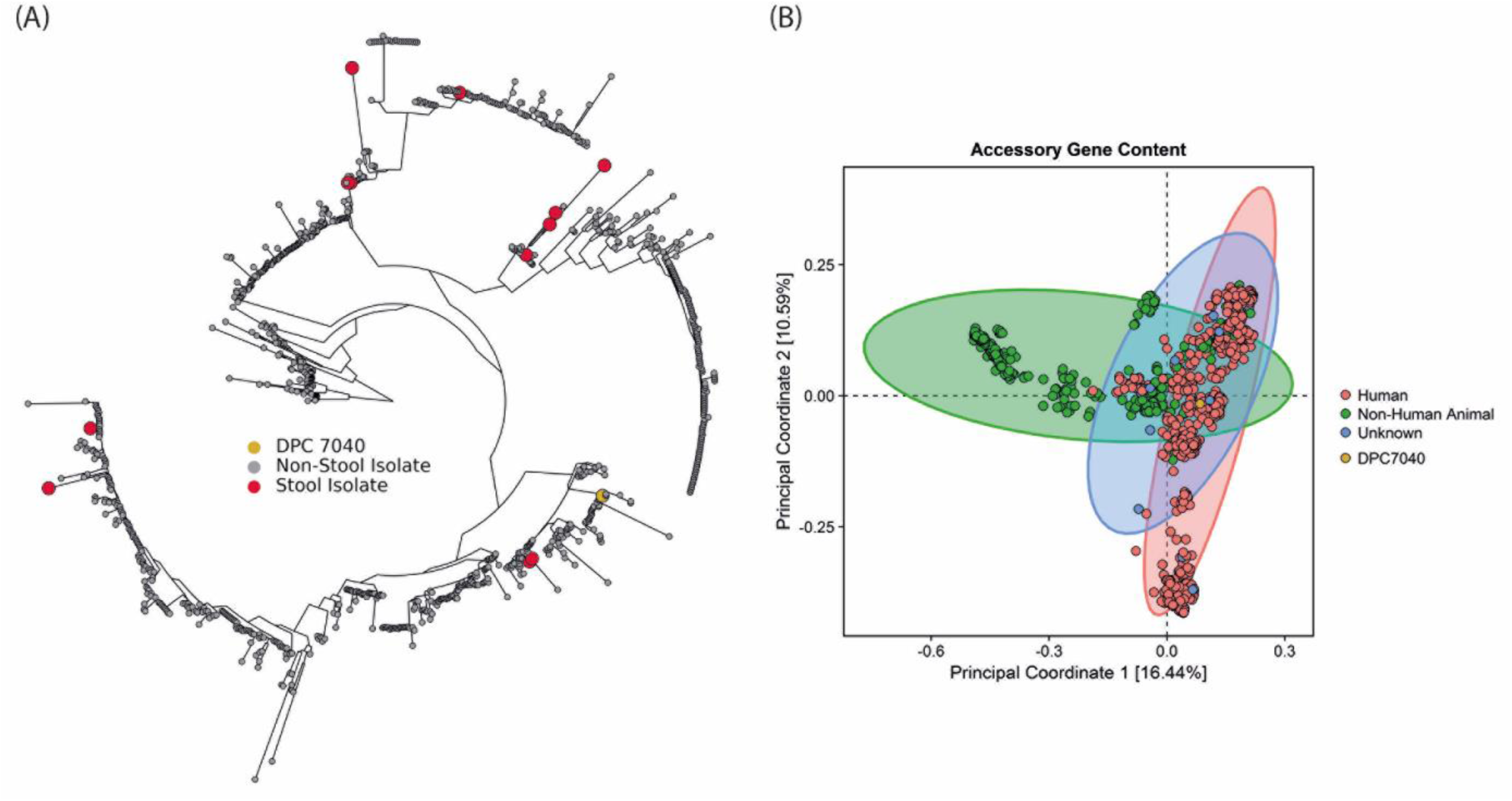
A. *S. agalactiae* phylogenetic tree of all genomes publicly available in PATRIC database 30/08/2019. The 12 stool isolates are marked in red and *S. agalactiae* DPC7040 is yellow. B. PCoA based on the presence/absence of accessory genes (genes present in less than 90% of the genomes) in the *S. agalactiae* pangenome.

### Characterization of the nisin P gene cluster from *S. agalactiae* DPC7040

The prototypical nisin A gene cluster consists of the structural gene *nisA*, followed by *nisBTCIPRKFEG* ^18^. The nisin P gene cluster in *S. agalactiae* DPC7040 has a similar gene order but the *nipPRKFEG* genes are located to the front of the *nipA* structural gene, a phenomenon which also occurs in the nisin U gene cluster ^19^ (Figure 3). The same gene order was observed when the nisin P gene cluster was first reported in the genome of *Streptococcus suis* ^20^.

**Figure 3.**
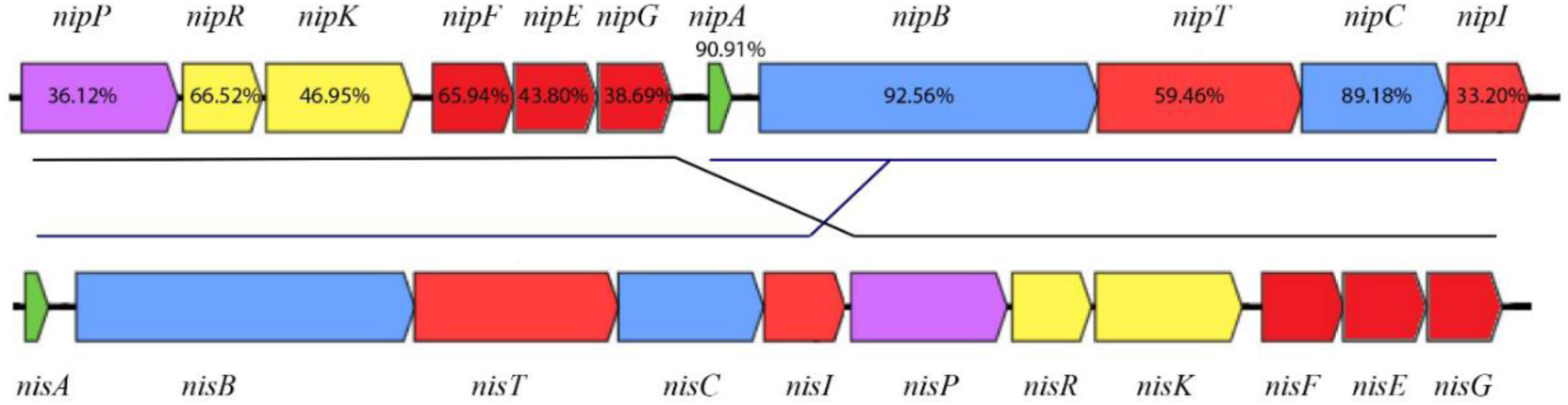
Representation of the bacteriocin-encoding nisin P gene cluster compared to the nisin A operon. Figures indicate percentage of amino acid similarity with nisin A homologues.

### Extraction and purification of nisin P

Nisin P and nisin H are produced by streptococci that require complex media (BHI and MRS, respectively) for growth and bacteriocin production. Nisin P production by *S. agalactiae* DPC7040 required initial induction by nisin A, taking advantage of the inducing activity of nisin peptides ^21^, but was subsequently self-inducing. The highest antimicrobial activity was achieved when *S. agalactiae* DPC7040 was cultured in BHI broth under anaerobic conditions for 48 h. Activity in well diffusion assays was less than nisin A (Figure 4A). Purification was achieved by a four-step process using Amberlite XAD16N, SP Sepharose, C18 SPE and Reversed phase HPLC. Overall yield was very low, i.e., 0.2-0.35 mg/l for nisin H and 0.1-0.2 mg/l for nisin P (Figure 4), while the yield for nisin A production is typically 3-4 mg/l. The lower yield obtained for nisin H and P could be partially due to less efficient purification, but it is most likely due to lower cell numbers, typically in the range of 10^8^ cfu/ml for *S. hyointestinalis* and *S. agalactiae* compared to 10^9^ cfu/ml for the nisin A producer, *L. lactis* NZ9000.

**Figure 4.**
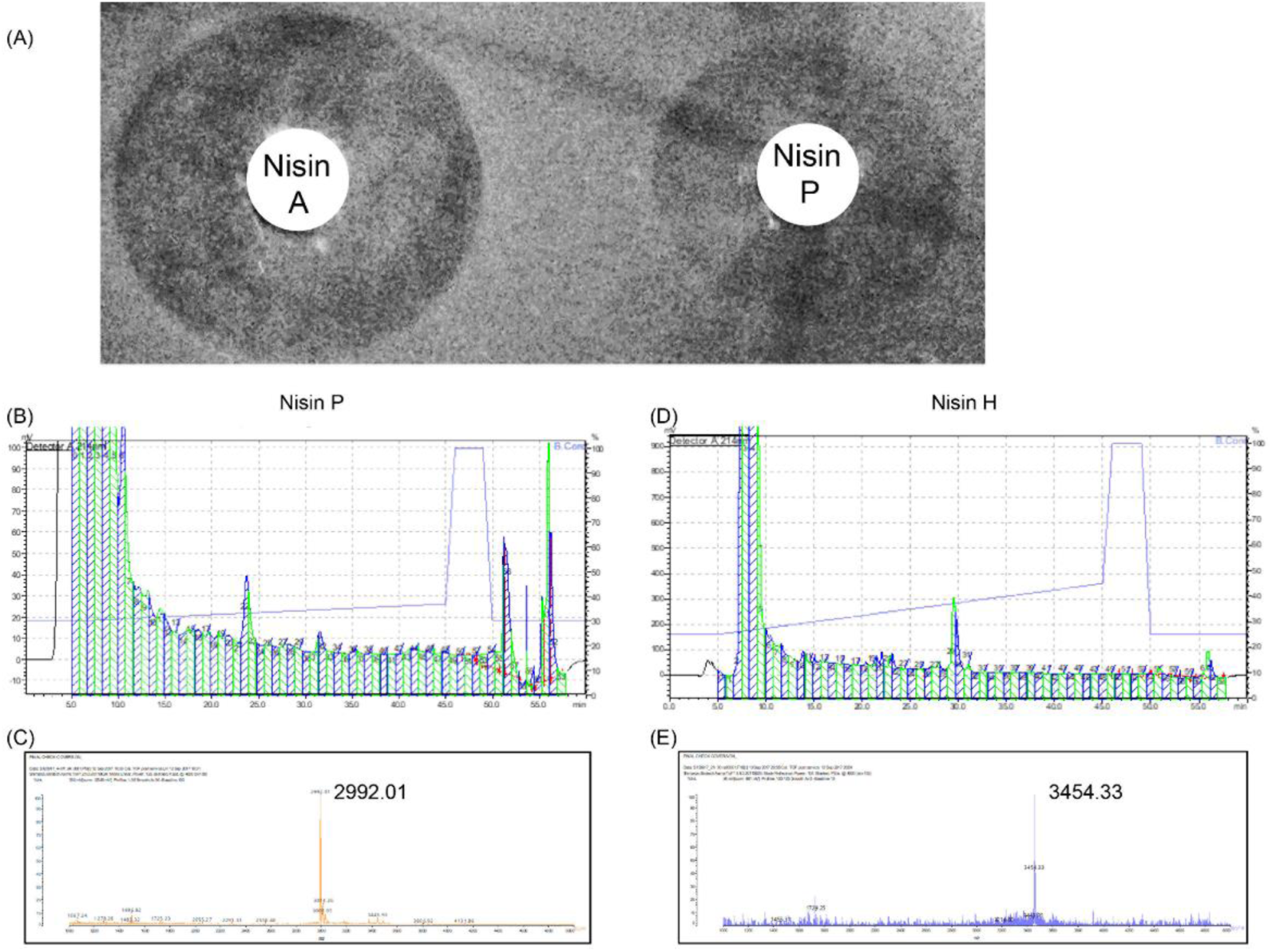
A. Inhibitory zones of nisin A and nisin P against indicator strain *L. bulgaricus* LMG 6901. B. Purification of nisin P from *S. agalactiae* DPC7040 and nisin H from *S. hyointestinalis* DPC6484 cultures. For nisin P: (B) RP-HPLC chromatogram and (C) MALDI TOF MS of active fraction number 24. Expected mass for nisin P is 2991 Da. For nisin H: (D) RP-HPLC chromatogram and (E) MS MALDI TOF MS of active fraction 30 showing mass corresponding to the expected nisin H mass of 3453 Da.

### Prediction of the nisin P structure

Purified nisin P peptide was tested to confirm that the expected peptide was intact. The extracted-ion chromatogram (XIC) shows the recovered nisin P at 9.56 min (Figure 5B). The base peak intensity chromatogram shows that some impurities were found (12.55 min), but the presence of intact nisin P was confirmed (Figure 5C).

**Figure 5.**
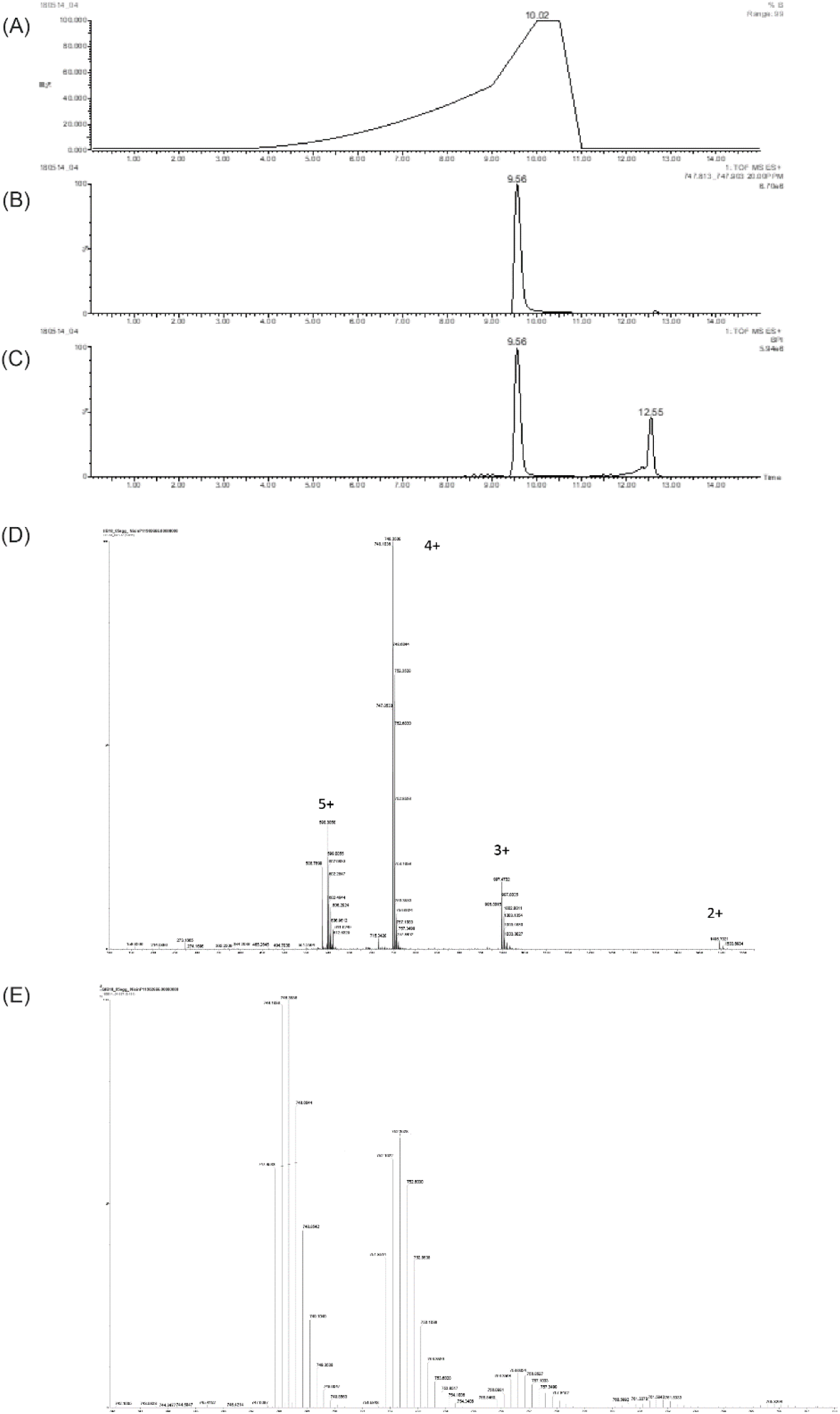
Chromatograms from LC-MS analysis of purified nisin P. A. Elution gradient of 1-50% acetonitrile in 0.1% formic acid in 9 min and to 100% acetonitrile in 1 min. B. Extracted-ion chromatogram (XIC) of nisin P. C. Base peak intensity chromatogram. D. Raw spectrum of series of charge states from 5+ to 2+. E. Spectrum detail of charge state 4+, confirming expected mass of purified nisin P. The mass calculation is performed by multiplying by 4 the 747.8533 ion mass and removing four charges of protons (747.8533*4) - (4*1.007276) = 2987.3841.

Possible modifications were further assessed by the analysis of the eluted samples, that showed a series of charge states ranging from 5+ to 2+ (Figure 5D). The most abundant charge state was 4+. A more detailed analysis of this isotope pattern showed a mass value of 2987.3841 Da, that matched the expected mass of fully dehydrated (−8 H_2_O) nisin P (2987.3922 Da) (Figure 5E). Additional peaks have an addition of 4 m/z corresponding to 16 Da, suggesting oxidation. Addition of 18 Da (H_2_O) (or multiples) cannot be seen, meaning that the major part of nisin P is completely dehydrated.

After trypsin digestion, 7 exclusive unique peptides with 30 exclusive unique spectra, matching the sequence of nisin P with 100% probability were detected (Table 2). Those spectra cover 31 of the 31 amino acid residues of nisin P corresponding to 100% sequence coverage.

**Table 2.** List of peptides obtained by digestion with trypsin and LC-MS/MS analysis.

| Domain | Tryptic peptide | m/z (1+) | MC | PSMs | CAM | Highest probability (%) |
| --- | --- | --- | --- | --- | --- | --- |
| N-term | VTSKSLCTPGCK | 1,207.58 | 1 | 342 | 145 | 100 |
| N-term | VTSKSLCTPGCKTGILMTCAIK | 2,162.09 | 2 | 6 | 2 | 7 |
| Core | TGILMTCAIK | 1,029.54 | 0 | 37 | 0 | - |
| Core | SLSLCTPGCKTGILMTCAIK | 1,820.91 | 1 | 4 | 0 | - |
| Full | VTSKSLCTPGCKTGILMTCAIKTATCGCHFG | 3,003.39 | 3 | 17 | 1 | 23 |
| C-term | TGILMTCAIKTATCGCHFG | 1,984.88 | 1 | 144 | 70 | 100 |
| C-term | TATCGCHFG | 916.3329 | 0 | 91 | 52 | 100 |

MC: Missed cleavage. PSMs: Peptide Spectra Match.

The presence or absence of rings can be determined by the suppression of the MS2 fragmentation within the regions that correspond to those rings. Two types of modifications were observed: dehydrations and carbamido-methylation from iodoacetamide alkylation of free cysteine residues (Figure 6). Dehydrations (−18) were present on S and T residues (Figure 6). Dehydrations on the T residue have been reported only in some nisin variants ^15, 22^. Carbamido-methylations (CAM, C+57) were observed on modified cysteine residues and are an indicator of the absence of a lanthionine ring, while unmodified cysteines indicate participation in ring formation. Several spectra from the complete undigested nisin P peptide appeared with no CAM modification and unfragmented except in the linear regions (Figure 6). This indicates the formation of all rings. Other spectra indicated incomplete ring formation. For example, the N-terminal peptide VTSKSLCTPGCK and the C-terminal peptides TGILMTCAIKTATCGCHFG and TATCGCHF showed CAM modifications in approximately 50% of the 342, 144 and 91 peptide spectra matches (PSMs) obtained by trypsin digestion with a probability of 100%. Another peptide, VTSKSLCTPGCKTGILMTCAIK and the full peptide, showed two modifications, with a low probability, in the same residues as VTSKSLCTPGCK. This indicates the absence of ring B in an estimated 50% of the peptides. The presence of CAM modifications in the C-terminus in peptides TGILMTCAIKTATCGCHFG and TATCGCHFG indicates the absence of ring E in an estimated 50% of the peptides. Based on these observations, the structure of nisin P was confirmed (Figure 7).

**Figure 6.**
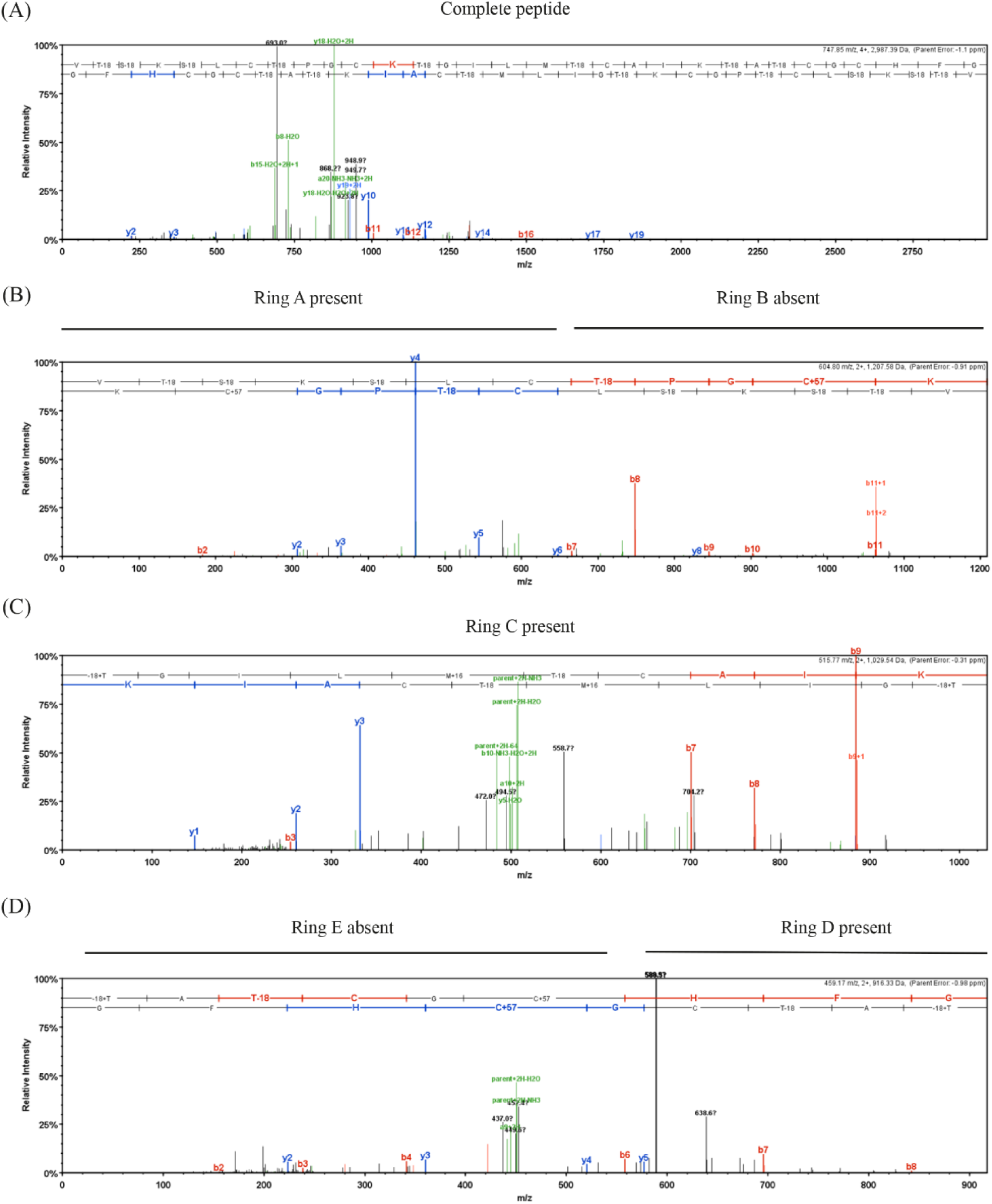
MS/MS spectra for nisin P peptides obtained by trypsin digestion and generated in Scaffold. b-ions are represented in red and y-ions in blue. A. Spectrum of nisin P showing fragmentation only in linear region, indicating presence of all rings. B. Spectrum showing presence of ring A (C is free and there is no fragmentation) and absence of ring B (C has a CAM modification and is fragmented). C. detail of presence of ring C and hinge region, AIK D. Presence of ring D and CAM modification indicating the absence of ring E.

**Figure 7.**
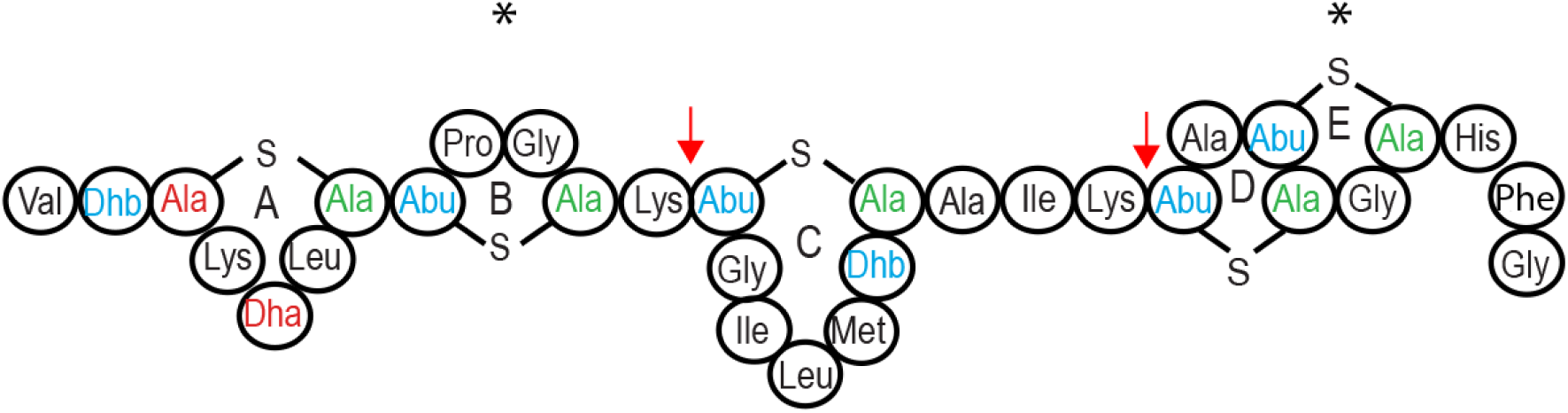
Fully modified structure of nisin P. Locations of the sites in nisin P cleaved by trypsin to form the different peptides summarized in table 2 are indicated by arrows, * indicates that the ring is not always formed.

### pH stability of nisin A, P and H

The antimicrobial activities of nisin A, P and H stored at pH 3, 5, 6, 7, and 8 for 24, 48 and 72 h were compared by well-diffusion assay using *L. bulgaricus* LMG 6901 as an indicator strain (Figure 8). Overall, nisin A antimicrobial activity was more stable than both P and H up to 72 h and at acidic pH. Nisin P was the least stable as it lost most inhibitory activity against *L. bulgaricus* LMG 6901 after 24 h and was inactive at 72 h at acidic pH, though it retained some activity at pHs 7 and 8. While nisin H activity was less stable than nisin A, it was more stable than nisin P as it retained more activity at 24 h, showing an optimum at pH 5, also at 48 and 72 h, and it showed some activity at 72 h at pHs 5, 6, 7 and 8.

**Figure 8.**
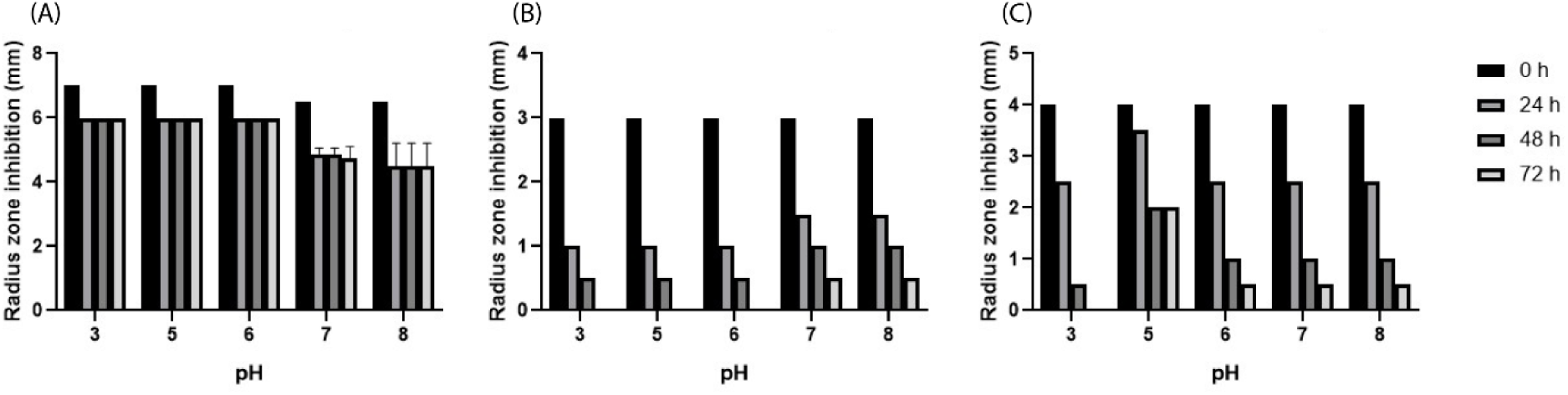
Stability of nisin A (A), P (B) and H (C) stored at different pH for 24, 48 and 72 h measured by radius (mm) of inhibitory activity against *L. bulgaricus* LMG 6901 using well-diffusion assay. Results are the mean of triplicate measurements +/-standard deviation.

### Cross-immunity assays

Purified nisin P, at 0.20 mg/ml, had inhibitory activity against natural nisin A and U producers but not against *S. hyointestinalis* DPC 6484 (nisin H producer, Table 3). A lower concentration of nisin A (0.022 µM) was required to inhibit *S. agalactiae* DPC 7040 than was required to inhibit *S. uberis* strain 42 (0.4 µM) and *L. lactis* NZ9800 (26µM). Nisin P inhibited *L. lactis* NZ9800 growth at 8.5 µM and *S. uberis* strain 42 at 4.25 µM. Nisin P inhibited the nisin P producer *S. agalactiae* DPC7040 at 17 µM. Nisin clusters encoding nisin A, U and P have two systems involved in immunity, one through the immunity protein NisI and the other one through the ABC transporter system NisFEG. However, the cluster for nisin H does not have a *nisI* gene ^15^ and yet shows immunity to nisins P and A. A Blastp analysis was performed on the NisFEG proteins from the four operons. The NisF- and NisG-determinants are similar in the nisin A, P and U operons, but NshF and NshG from the nisin H cluster have a very low percentage of similarity to equivalent proteins from the nisin A, P and U gene clusters (13.8%, 14.1% and 15.4% to NshF and 9%, 8.9% and 9.8% to NshG, respectively). NshE and NisE showed 48.3% and 43.8% identity to NipE, while NsuE was more similar to NipE (79.8% identity).

**Table 3.** Summary of inhibitory activity of nisin P and A against different natural nisin producers

| Strain | Activity nisin P | Activity nisin A | MIC nisin P | MIC nisin A |
| --- | --- | --- | --- | --- |
| *L. lactis*<br>NZ9800 (nisin A) | ++ | + | 8.5 µM | 26 µM |
| *S. uberis*<br>strain 42 (nisin U) | ++ | +++ | 4.25 µM | 0.4 µM |
| *S. hyointestinalis*<br>DPC6484 (nisin H) | - | - | - | - |
| *S. agalactiae*<br>DPC 7040 (nisin P) | + | +++ | 17 µM | 0.022 µM |

-, No activity; +, 1 mm radius inhibition zone; ++, 1-5mm radius inhibition zone; +++, >5mm inhibition zone.

### Spectra of inhibition and MIC of purified nisin A and nisin P peptides against a range of strains

In general, purified nisin A was more potent and had a wider spectrum of inhibition than nisin P (Table 4). Nisin P inhibited all lactobacilli and staphylococci assayed, but inhibition was at higher concentrations than that observed for nisin A, which was also able to inhibit the growth of *Listeria* and enterococci. This is consistent with the spectrum of action of other nisins, that are generally potent against lactobacilli and staphylococci, and, indeed, Gram positive bacteria in general ^23^.

**Table 4.** Summary of inhibitory activity of nisin P and A

| Strain | Activity<br>nisin P | Activity<br>nisin A | MIC<br>nisin P | MIC<br>nisin A |
| --- | --- | --- | --- | --- |
| *S. enterica* LT2 | - | - | - | - |
| *E. coli* ATCC 25922 | - | - | - | - |
| *C. sakazakii* NCTC 11467 | - | - | - | - |
| *C. perfringens* NCTC 3110<sup>a</sup> | - | - | - | - |
| *L. innocua* NCTC 11288 | - | - | - | - |
| *L. bulgaricus* LMG 6901 | +++ | +++ | 0.01 µM | 0.0005 µM |
| *L. monocytogenes* 3564 | - | ++ | - | - |
| *E. faecium* 4955 | ++ | +++ | 0.265 µM | 0.1 µM |
| *E. faecalis* LM1 | - | ++ | - | 6.5 µM |
| *E. faecalis* LM4 | - | ++ | - | 6.5 µM |
| *P. aeruginosa* 2064 APC | - | - | - | - |
| *S. aureus* 7016 | ++ | +++ | 0.531 µM | 0.05 µM |
| *S. capitis* 20G | ++ | +++ | 8.5 µM | 0.812 µM |
| *S. simulans* 2B10 | ++ | +++ | 0.531 µM | 0.325 µM |
| *S. epidermidis* 99 | + | +++ | 34 µM | 0.812 µM |
| *L. gallinarum* Lac43 | + | ++ | 8.5 µM | 0.812 µM |
| *L. frumenti* Lac53 | +++ | +++ | 1.06 µM | 0.27 µM |
| *L. reuteri* Lac54 | + | ++ | 8.5 µM | 0.812 µM |
| *L. paracasei* LacR1 | ++ | +++ | 17 µM | 1.62 µM |
| *L. plantarum* Lac 29 | +++ | +++ | 1.06 µM | 0.1 µM |
| *L. mucosae* Lac28 | +++ | +++ | 1.06 µM | 0.54 µM |
| *L. reuteri* Lac 56 | + | ++ | 69 μM | 3.25 μM |

<sup>a</sup> *C. perfringens* NCTC 3110 has usually been reported to be sensitive to nisin A
-, No activity; +, 1mm radius inhibition zone; ++, 1-5mm radius inhibition zone; +++, >5mm
inhibition zone.

### Nisin activity in fecal fermentation

The analysis of total live bacteria in faecal samples after fermentation in the presence of two concentrations (15 µM and 50 µM) of nisin A, P and H showed different consequences depending on nisin type and treatment (Figure 9). There were significant differences in the total numbers of live cells between the control and the treatments with nisin A, P and H, both at 8 and 24 h. Treatment with nisin A showed significantly less live cells compared to the controls at 8 h, showing Log_10_ 4.234 ± 0.023 and Log_10_ 4.792 ± 0.01 at 15 µM and 50 µM respectively, compared to Log_10_ 5.22 ± 0.005 in the control. At 24 h, after single nisin A treatments, bacterial regrowth was evident. The biggest impact was seen in samples which received a second nisin A treatment at 8 h, which showed Log_10_ 4.592 ± 0.022 and Log_10_ 5.248 ± 0.01 of live cells at 15 µM and 50 µM at 24 h compared to Log_10_ 5.22 ± 0.005 and Log_10_ 5.461± 0.019 in the control. Treatments with nisin P and nisin H showed higher numbers of live cells with all treatments, the numbers of live cells being closer to those of total cells.

**Figure 9.**
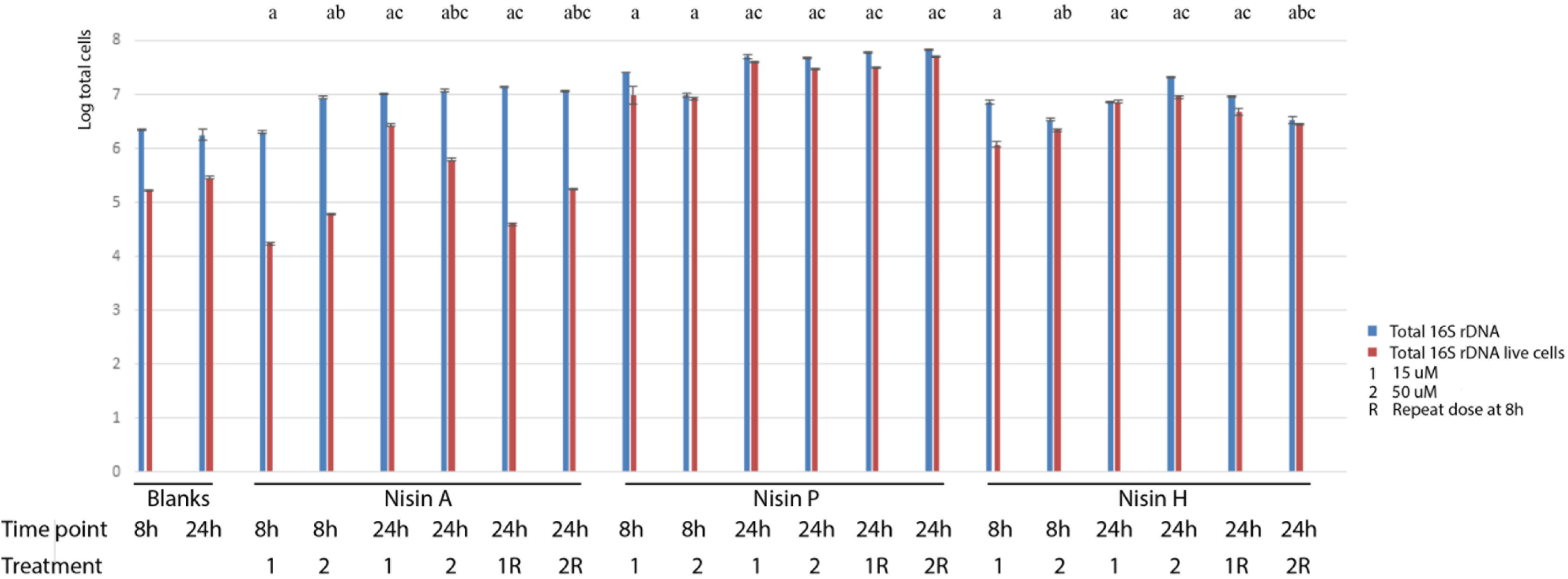
Total bacteria after nisins A, P and H treatments measured by 16S rDNA qPCR after fecal fermentation at 8 and 24 h. Two concentrations of peptide were tested, 15 µM and 50 µM. Half of the wells received a second dose of the same concentration at 8 h. Results are the mean of duplicate measurements +/-standard deviation. Statistics was conducted with live cells data: a, indicates significant difference between the treatment and the blank at 8 or 24 h; b, indicates significant difference between 15 µM and 50 µM concentrations; c, indicates significant difference between 8 and 24 h within the same nisin treatment (p<0.05).

### Activity induction

Nisin peptides also serve as auto-inducers of nisin production. The ability of nisin P, A and H to activate the nisin A promotor, fused to a *gfp* reporter, was assessed and found to differ (Table 5). The promoter was more sensitive to nisin A (1 ng/ml – 1 µg/ml) than nisin H (10 ng/ml – 1 µg/ml) and nisin P (100 ng/ml – 10 µg/ml). Higher concentrations of nisin P were required to activate the promotor, but it continued to induce promoter activity at higher concentrations (10 µg/ml) whereas nisin A and H were capable of inducing the promoter only up to 1 µg/ml concentrations of peptides.

**Table 5.** Fluorescence levels of nisin induction of *L. lactis* NZ9000 pNZ8150 *gfp*+ by nisin P, A and H.

|  | 1<br>mg/ml | 10 ug/ml | 1 ug/ml | 500<br>ng/ml | 100<br>ng/ml | 50<br>ng/ml | 20<br>ng/ml | 10<br>ng/ml | 5 ng/ml | 2 ng/ml | 1<br>ng/ml |
| --- | --- | --- | --- | --- | --- | --- | --- | --- | --- | --- | --- |
| Nisin<br>P | - | 1758.33<br>± 41.66 | 658.33 ±<br>8.33 | 316.66 ±<br>33.33 | 75 ±<br>8.333 | - | - | - | - | - | - |
| Nisin<br>A | - | - | 1408.33<br>± 25 | 1916.5 ±<br>83.5 | 1458.33<br>± 25 | 1050 ±<br>0 | 549.99<br>± 16.66 | 358.33<br>± 8.33 | 224.99±<br>8.33 | 116.66<br>± 16.66 | 83.33<br>±<br>16.66 |
| Nisin<br>H | - | - | 1566.66<br>± 33.33 | 1133.33<br>± 66.66 | 608.33<br>± 41.66 | 358.33<br>± 8.33 | 208.33<br>± 8.33 | 108.33<br>± 25 | - | - | - |

## Discussion

This is the first report of production of nisin P, a natural variant of nisin, originally predicted on the basis *in silico* of genomic analysis of other two streptococcal species ^20, 24^. Initially, genes related to nisin P production were identified by phylogenetic categorization of LanB- and LanC-encoding enzymes in *Streptococcus pasteurianus*, and analysis of the associated *lanA* gene suggested the ability of this strain to produce an analogue of nisin U that was designated nisin P ^24^. The gene cluster was later reported in *Streptococcus suis* ^20^. The fecal isolate that was the focus of this study, *S. agalactiae* DPC7040, contained an intact nisin P cluster and produced nisin P after initial induction with nisin A.

The genome of *S. agalactiae* DPC7040 was compared with 993 *S. agalactiae* genomes publicly available in the PATRIC database. This showed that *S. agalactiae* did not cluster phylogenetically by human body isolation site. However, PCoA showed different clustering between all strains of human origin and the strains of non-human animal origin based on accessory gene content. No unique genome features could be associated with *S. agalactiae* DPC7040, apart from the presence of the nisin P lantibiotic cluster, which was absent from the other 12 stool isolates.

When nisin P activity was compared to nisin A and H, a lower inhibitory effect was observed. Nisin P showed a higher MIC against a panel of food and gut isolates, and also had less effect on the viability of the microbiota in a fermentation of human fecal material, despite being sourced from a human gut isolate. One consideration is differences in stability, as both nisin H and particularly nisin P lost a significant amount of activity after 24 h at pH 6-7. The differences in bioactivity are also likely to be contributed to by structural differences between the molecules. Nisin P is three amino acids shorter than nisin A at the important pore-forming C-terminal domain. Previously, removal of amino acids from the C-terminal region of nisin A, affecting rings D and E, showed a decrease of up to 100 fold of its inhibitory activity against *L. lactis* and 50 fold against *Micrococcus luteus*, suggesting that different conformations might have a different outcome against different bacteria ^25–27^. The N-terminal part of the nisin A molecule, that includes rings A, B and C, has been shown to be responsible for the binding of nisin to the membrane surface and / or its oligomerization ^25^. Ring B is known for being very sensitive in substitution studies and the removal of ring B in nisin A also gave a loss of activity in nisin mutants ^28^, so the absence of rings B and E in 50% of the nisin P peptides may contribute to reduced activity.

Another important region involved in nisin bioactivity is the hinge region, at positions 21-23 ^29, 30^. The hinge region consists of the three amino acids (NMK in nisin A, AIK in nisin P) that are responsible for reorientation after binding with lipid II to penetrate into the target cell membrane ^31^ and importance in nisin antimicrobial activity has been established through bioengineering studies ^30–33^. These studies suggest a preference for small, chiral amino acids in that region ^32^. Nisin H has an aromatic polar tyrosine instead of the non-polar methionine in the second position. Nisin P has two changes in the hinge region, with an exchange of a polar asparagine for a non-polar alanine, and an exchange of methionine with an isoleucine, both non-polar amino acids. Directed amino acid changes in nisin A showed that alanine in position 21 reduced bioactivity against methicillin resistant *Staphylococcus aureus* ST528, *S. aureus* DPC5245 and *S. agalactiae* ATCC13813 to half of the bioactivity from nisin A; however, molecules carrying an isoleucine in position 22 showed an enhanced bioactivity against the three microorganisms ^33^. Nisin P might also receive a benefit from the presence of a lysine residue in position 4 instead of isoleucine, as previous studies have shown that the presence of an additional positive charge in ring A has a positive effect on activity ^28^. Further studies are required to determine if there are indeed instances where nisin P benefits by virtue of these changes.

Cross-immunity assays indicated that nisin P was able to inhibit the growth of the nisin A and nisin U producers, suggesting that their immunity systems are so different as to not provide protection from the effect of nisin P. *S. agalactiae* DPC7040 was also very sensitive to nisin A. Nisin P and nisin A inhibited the growth of the nisin U producer, *S. uberis* strain 42. Although the impact of nisin A against this strain has been reported previously ^15^, it contrasts with initial reports that *S. uberis* strain 42 was cross-resistant to nisin A ^13^. Nisin A and nisin P were not able to inhibit the growth of *S. hyointestinalis* DPC6484, the nisin H producer, an observation previously made in the case of nisin A ^15^. *S. hyointestinalis* DPC6484 does not have one of the two systems involved in lantibiotic immunity, NisI. NisI has been reported to have a more crucial role in immunity than the NisFEG ABC transporter system, which was considered to provide around 20% of the wildtype immunity ^34–36^. However, another study highlighted that *nisI* expression provided a very low level of immunity (1-4%) ^37^. Regardless, the immunity of *S. hyointestinalis* DPC6484 against other nisin peptides despite the low levels of similarity of NshFEG to the homologues from the nisin A, P and U producers is somewhat surprising.

From an autoinduction perspective, in addition to having a lower antimicrobial activity, nisin P also required higher concentrations to activate the nisin A promoter. However, this might be due to the specificity of the molecule or the receptor. Given the structural differences between the nisin variants of lactococcal and streptococcal origin, it would be interesting to investigate if using different nisin inducers with cognate receptors impacts on levels of induction.

In conclusion, the study of both bioengineered and natural variants of nisin aids the understanding of structure-activity relationships of this molecule and that knowledge can be applied to other lantibiotics and to the further development of more efficient antimicrobials. Nature provides a nisin variation system and previously unknown nisin variants continue to be discovered. The wide dissemination of the clusters encoding this molecule could be linked to its ability to provide a competitive advantage and increased fitness. Future investigation of nisin P modification and production in mixed communities might help to establish further knowledge on whether it would be a good candidate as an antimicrobial agent in certain environments.

## Materials and methods

### Bacterial strains, media and culture conditions

Bacteria were sourced from culture collections (ATCC, American Type Culture Collection, NCTC, National Collection of Type Cultures) and in-house collections at Teagasc and Quadram Institute Bioscience. *L. lactis* NZ 9800 (nisin A producer) and *L. lactis* NZ 9000 pNZ8150 *gfp*+ DPC7179 were grown aerobically at 30°C in GM17 (Oxoid - CM0817). All other bacteria were cultured at 37°C. *S. uberis* strain 42 (nisin U producer), *S. hyointestinalis* DPC6484 (nisin H producer) and *S. agalactiae* DPC7040 (nisin P producer) were grown in BHI (Merck – 110493) anaerobically, as were indicator strains *Cronobacter sakazakii* NCTC 11467, *Clostridium perfringens* NCTC 3110 and *Listeria innocua* NCTC 11288. *Listeria monocytogenes* 3564, *Pseudomonas aeruginosa* 2064 APC, *Staphylococcus aureus* 7016, *Staphylococcus capitis* 20G, *Staphylococcus simulans* 2B10 and *Staphylococcus epidermidis* 99 were grown aerobically in BHI; *Salmonella enterica* LT2 and *Escherichia coli* ATCC 25922 were grown anaerobically in LB (Merck – 110285); *Lactobacillus bulgaricus* LMG 6901, *Enterococcus faecalis* LM1, *E. faecalis* LM4, *Lactobacillus gallinarum* Lac 43, *Lactobacillus frumenti* Lac 53, *Lactobacillus reuteri* Lac 54, *L. reuteri* Lac 56, *Lactobacillus paracasei* Lac R1, *Lactobacillus plantarum* Lac 29 and *Lactobacillus mucosae* Lac 28 were grown anaerobically in MRS and *Enterococcus faecium* 4955 was grown in MRS (Oxoid - CM0361) aerobically. Anaerobic growth was performed in an anaerobic chamber (Don Whitley, UK) in an atmosphere of 5% CO_2_, 10% H_2_ and 85% N_2_.

### *Genomic analysis of* S. agalactiae *DPC7040*

*S. agalactiae* DPC7040 was isolated from a human fecal sample of a male donor with body mass index of less than 25. Initial culture was performed anaerobically at 37 °C for 48 h using Wilkins-Chalgren anaerobe agar (Oxoid - CM0643). A genomic library was prepared from gDNA using the Nextera XT DNA library preparation kit (Illumina, USA). Whole genome sequencing was performed using Illumina’s MiSeq platform with the MiSeq V3 600 cycles Paired Ends kit using paired-end 2×300 bp reads. The quality reads examination and the trimming of low-quality bases and Illumina adaptors were conducted using FastQC and Trim-Galore ^38^. Assembly of contigs from paired-end reads was achieved using SPAdes ^39^. Bacteriocin operons were identified using BAGEL3 ^40^ and manually curated using Artemis ^41^. Accession number for *S. agalactiae* DPC7040 is WIDP00000000. The *S. agalactiae* pan-genome was constructed by downloading all available genomes labelled “*Streptococcus agalactiae*” from the PATRIC database (30/08/2019). The genomes identified as non-*S. agalactiae* using Kraken2 (v. 2.0.7) and PhyloPhlAn (v. 0.99) ^42^ were removed from the analysis, leaving 993 genomes that were annotated using Prokka (v.1.11). A pangenome was constructed using Roary (v. 3.12.0) ^43^ using default parameters, and Scoary ^44^ was used to perform pangenome-wide associations for stool vs. non-stool, and to look for genes significantly associated with stool isolates. Blastp ^45^ was used to manually annotate genes with significant associations using the pangenome reference sequences produced by Roary. A phylogenetic tree was built using PhyloPhlAn (v. 0.99) ^42^ and visualized using GraPhlAn ^46^. Principal Coordinates analyses (PCoA) were performed using the presence/absence output from Roary analysis, converted into binary format in R (v. 3.4.4) with removal of genes which were present in over 90% of the genomes to focus on the accessory genome of the species. The principal coordinate model was built based on Jaccard distance between each accessory genome using the ‘vegan’ package (v. 2.5-1) ^47^ and visualized using ggplot2 (v. 2.2.1) ^48^. Additionally, LanB and LanC were searched manually in genomes of the 12 stool isolates *S. agalactiae* GBS10, 11 and 12 and MC625 to MC629 and MC631 to MC634.

### Purification of nisin A, H and P

Nisin P production required initial induction with 500 ng/ml of nisin A, which was not included in subsequent *S. agalactiae* DPC 7040 subcultures. Nisin P was purified from 2 l of *S. agalactiae* DPC 7040 grown anaerobically in BHI for 48 h (2.0 × 10^8^ cfu/ml). Supernatant was obtained by centrifuging the culture at 8000 x g, 10 min. Culture supernatant was applied to a 30 ml SP Sepharose column pre-equilibrated with 20 mM sodium acetate pH 4.4. The column was washed with 100 ml of 20 mM sodium acetate pH 4.4 and then eluted with 150 ml of 20 mM sodium phosphate buffer pH 7 containing 1 M NaCl. The NaCl-containing eluate was applied to a 60 ml, 10 mg Strata–E C18 SPE column (Phenomenex, UK) pre-equilibrated with methanol and water. The column was washed with 60 ml 30% ethanol and antimicrobial activity eluted with 70% 2-propan-ol 0.1% TFA (IPA). Antimicrobial activity of eluents from purification steps were tested via well-diffusion assay: indicator plates were prepared by adding 100 µl of an overnight culture of the indicator strain to 20 ml of melted agar, then 50 µl of each test solution was added to wells made in the agar and plates were incubated overnight at 37°C using the conditions required by the indicator strain.

The IPA was removed from the C18 SPE eluent by rotary evaporation and sample applied to a semi preparative Proteo Jupiter (10 x 250 mm, 90Å, 4µ) RP-HPLC column (Phenomenex, Cheshire, UK) running a 30-37% acetonitrile 0.1% TFA gradient where buffer A was 0.1% TFA and buffer B was 100% acetonitrile, 0.1% TFA. The eluate was monitored at 214 nm and fractions were collected at intervals. Nisin P-containing fractions deemed pure by Matrix-Assisted Laser Desorption/Ionization-Time Of Flight (MALDI-TOF) mass spectrometry (MS) were lyophilized using a Genevac lyophilized (Suffolk, UK).

Nisin H was purified from 2 l of culture (1.2 × 10^8^ cfu/ml) grown overnight at 37°C in TSB (Oxoid) without induction and purified as for nisin P except a 25-40% acetonitrile 0.1% TFA gradient was used for HPLC purification. Nisin A was purified from nisin A supplied by Handary SA (Belgium) by reversed phase HPLC using a 25-45% acetonitrile 0.1% TFA gradient as described above.

Nisin P, H and A HPLC fractions containing antimicrobial activity against *L. bulgaricus* LMG 6901 were analyzed via MALDI-TOF MS to confirm the molecular mass of the antimicrobial peptide and to assess peptide purity. HPLC fractions deemed pure by MALDI-TOF MS were combined and lyophilized in a Genevac lyophilizer. MALDI-TOF MS was performed with an Axima TOF^2^ MALDI-TOF mass spectrometer in positive-ion reflectron mode (Shimadzu Biotech, UK).

### Nisin P structural analysis

The intact peptide was analysed by LC-MS on a Synapt G2-Si mass spectrometer coupled to an Acquity UPLC system (Waters, UK). Aliquots of the sample were injected onto an Acquity UPLC® BEH C18 column, 1.7 µm, 1×100 mm (Waters, UK) and eluted with a gradient of 1-50% acetonitrile in 0.1% formic acid in 9 min and then ramped to 100% acetonitrile in 1 min at a flow rate of 0.08 ml/min with a column temperature of 45°C.

The LC-MS was operated in positive MS-TOF resolution mode and with a capillary voltage of 3 kV and a cone voltage of 40 V in the range of m/z 100–2000. Leu-enkephalin peptide 0.5 µM (Waters −186006013) was infused at 10 µl/min as a lock mass and measured every 20 s. Masslynx 4.1 software (Waters, UK) was used to generate the spectra by combining several scans. The peaks were centered, and the mass calculated from the monoisotopic peak.

For structure confirmation, purified nisin P was digested with trypsin (Sigma) and analyzed using nanoLC-MS/MS on an Orbitrap Fusion™ Tribrid™ Mass Spectrometer coupled to an UltiMate® 3000 RSLCnano LC system (Thermo Scientific, UK). To investigate ring formation, samples were treated with iodoacetamide to alkylate free cysteine residues. The samples were loaded and trapped using a pre-column which was then switched in-line to the analytical column for separation. Peptides were separated on a nanoEase *m/z* column (HSS C18 T3, 100 Å, 1.8 µm; Waters, UK) using a gradient of acetonitrile at a flow rate of 0.25 µl/min with the following solvent steps A (water, 0.1% formic acid) and B (80% acetonitrile, 0.1% formic acid): 0-4 min 3% B (trap only); 4-15 min increase to 13% B; 15-77 min increase to 38% B; 77-92 min increase to 55% B; followed by a ramp to 99% B and re-equilibration to 3% B.

Data dependent analysis was performed using parallel CID and HCD fragmentation with the following parameters: positive ion mode, orbitrap MS resolution = 60k, mass range (quadrupole) = 300-1800 m/z, MS2 top20 in ion trap, threshold 1.9e4, isolation window 1.6 Da, charge states 2-5, AGC target 1.9e4, max inject time 35 ms, dynamic exclusion 1 count, 15 s exclusion, exclusion mass window ±5 ppm. MS scans were saved in profile mode and MS2 scans were saved in centroid mode.

MaxQuant 1.6.2.3 was used to generate recalibrated peaklists, and the database search was performed with the merged HCD and CID peak lists using Mascot 2.4.1 (Matrixscience, UK). The search was performed, with a precursor tolerance of 6 ppm and a fragment tolerance of 0.6 Da, on a *S. agalactiae* protein sequence database, which included the nisin P gene cluster (Uniprot, January 2018, 2,123 sequences) to which the nisin P peptide sequence had been added. The enzyme was set to trypsin/P with a maximum of 2 allowed missed cleavages (MC). Dehydration (−18 Da) of serine and threonine, oxidation of methionine and carbamido-methylation (CAM, +57 Da) of cysteine were set as variable modifications. The Mascot search results were imported into Scaffold 4.4.1.1 (www.proteomsoftware.com) using identification probabilities of 99% for proteins and 95% or 0% for peptides, as discussed in the results.

### pH stability of nisin P compared to nisin H and A

Nisin P, A and H were resuspended at 1 µg/ml in distilled water. Duplicate samples at pHs 3, 5, 6, 7 and 8 were prepared for each nisin. Antimicrobial activity was tested at 24, 48 and 72 h via well-diffusion assay using *L. bulgaricus* LMG 6901 as indicator.

### Cross-immunity assays

Cross immunity assays were conducted by well diffusion assay to test the activity of the purified nisin P and nisin A against natural nisin A, P, H and U producers. 100 μl of an overnight inoculum of each indicator strain was added to 20 ml of melted agar media. 50 μl of pure peptides at 0.20 mg/ml were added to a well pre-formed in the agar and incubated overnight according to the growth requirements of the different nisin producers used as indicator strains. Growth of the indicator strain in the presence of peptide was taken as confirmation of immunity. Blastp analysis ^45^ was performed on the proteins responsible for the immunity system, NisF, functioning as an ATPase, and NisE and NisG, the membrane spanning domains of the transporter, of natural producers of nisin A, P, H and U.

### Spectrum of inhibitory activity

The inhibitory activity of nisin P compared with nisin A against a range of indicator strains was determined by well diffusion assay. 50 µl of the purified peptides at a concentration of 0.2 mg/ml were added to wells preformed in the indicator plates, which were incubated overnight according to growth requirements of the indicator strain. Inhibitory activity was determined by the presence of a zone of inhibition.

### Minimum inhibitory concentration (MIC) assays

96-well plates (Sarstedt, UK) were pre-treated with bovine serum albumin (BSA) on the day of assay. For this purpose, 0.5 g of BSA (Sigma, UK) was mixed with 2.5 ml of 20X phosphate buffered saline (PBS, Sigma, UK). Water was added to the mix for a total volume of 50 ml. 200 µl were added to each well and the plate was incubated for 30 to 45 mins at 37°C. After this time, BSA was removed and the wells washed with 1X PBS and air dried in a sterile hood. For MICs, peptide concentrations of 4X the highest test concentration were prepared in 350 µl of the growth media required for the target strain to grow, and 100 µl of this mix was added to the first well of each row. 100 µl of the same media were added to each well and 2-fold serial dilution was performed alongside the row. After this, 100 µl of culture inoculated with the target strain was added to each well. To prepare this inoculum, an overnight culture was sub-cultured the day of the assay and incubated until an optical density (OD_600_) of 0.5 was reached. The culture was diluted 1:10 and 20 µl of this added to 980 µl of media. 150 µl of this new mix was added to 14.85 ml of growth media. The objective was to standardize the final inoculum to 10^5^ cfu/ml in 200 µl. The plate was incubated under the requirements of the target strain, and MIC determined as the lowest peptide concentration where there was no growth of the target strain. OD_600_ was measured using a Synergy HT Microplate reader (BioTek, UK).

### MicroMatrix fermentations

#### Preparation of a fecal standard

A fecal standard for fermentation in a micro-Matrix was prepared following methodology described by O’Donnell et al ^49^. Fecal samples were provided by different donors, from a study approved by the Clinical Research Ethics Committee of the Cork Teaching Hospitals, UCC, (ECM 4 (k) 04/12/18). The samples of six healthy donors with no antibiotic treatment in the previous six months were collected and maintained at 4°C for 1-2 h before being transferred to an anaerobic chamber (5% CO_2_, 10% H_2_ and 85% N_2_) for processing. 200 g of feces were placed in a Circulator 400 stomacher bag (Seward, UK) with 200 ml of PBS containing 0.05% (w/v) cysteine hydrochloride. After filtering and homogenizing, the fecal slurry was centrifuged (4000 x g, 25 min) and resuspended in PBS. Sterile glycerol was added to the mix to a final concentration of 25% and aliquots were stored at −80°C until required.

#### MicroMatrix fermentation

Fecal standards were defrosted at 37 °C before use and prepared at 10% concentration in fecal medium as described by Fooks and Gibson ^50^. 2 ml of the mix were added to each well of a MicroMatrix 24-well cassette (Applikon ® Biotechnology, Netherlands). Purified nisin A, P and H were added to the wells to final concentrations of 15 or 50 µM. Fermentations took place for 24 h at 37°C, with pH of the medium adjusted and maintained at 6.8 in a MicroMatrix fermenter (Applikon ® Biotechnology). At time point 8 h, half of the nisin-treated wells received an additional dose of nisin A, P and H at the same concentration as at time 0 h. 500 μl aliquots were withdrawn from each well at time points 0, 8, 24 h (T0, T8, T24) and kept frozen for further analysis.

#### qPCR

200 μl aliquots from each MicroMatrix fermentation were taken for DNA extraction using QIAmp Fast DNA Stool Mini Kit (Qiagen, Crawley, UK), following the manufacturer’s instructions. To compare total cell numbers with live cells, ethidium bromide monoazide (EMA, Sigma, UK) treatment was performed by adding EMA to the sample at a concentration of 10 µg/ml, incubating the samples in darkness for 5 min and exposing them to 70-Watt HQI light for 10 min at 20 cm while keeping the samples on ice ^51^. Samples were then centrifuged at 12000 x g for 5 min and the supernatant discarded before proceeding with the standard DNA extraction with QIAmp Fast DNA Stool Mini Kit. Absolute quantification by qPCR was performed to determine bacterial numbers, using the Roche LightCycler 480 II platform. Although bacteria can have multiple copies of rDNA ^52^, we used the amplification of this gene as a reference to measure the effect of the different treatments. To quantify 16S bacterial counts, a standard curve was created using 1000 to 100 copies of 16S rDNA/μl. For amplification, 16S rDNA primers used were CO1 forward primer (5′-AGTTTGATCCTGGCTCAG-3′), and CO2 reverse primer (5′-TACCTTGTTACGACT-3′)^53^ with KAPA Lightcycler 480 mix (KAPA Biosystems Ltd., UK) according to manufacturer’s instructions. All samples were run in triplicate.

### Statistical analysis

Significant differences between groups were established using a paired t-test, assuming normal distribution, equal variances. Both sides of the distribution were considered. Significance was considered when P value was <0.05. Calculations were performed using Excel 365.

### Activity induction

Induction experiments were performed using *L. lactis* NZ9000 pNZ8150 gfp+, where gfp acts as a reporter of expression from a nisin A inducible promoter ^54^. An overnight culture of this strain was diluted 1:100 into fresh media and incubated until the OD_600_ reached 0.5. Purified nisin P, A and H were added to 1 ml of GM17 at a concentration of 1 mg/ml, 10 μg/ml, 1 μg/ml, 500, 100, 20, 10, 5, 2 and 1 ng/ml and 20 µl of the *L. lactis* NZ9000 pNZ8150 gfp+ were added to the mix. 200 µl were transferred to black 96-well plates (Nunc^TM^, Thermo Fisher, UK) and GFP was detected using a Synergy HT Microplate reader (BioTek, UK). The excitation filter was set at 485 nm and the emission filter at 528 nm.

## Acknowledgements

The authors would like to thank Elaine Lawton, Dr. Taís Kuniyoshi, Dr. Michelle O’Donnell and Dr. Silvia Arboleya for training and assistance.

