## Supplementary material for "First evidence of production of the lantibiotic nisin P": Table S1

| **Table S1.** Genes significantly associated with Streptococcus agalactiae stool isolates by presence or absence. | | | | | | | | | | | | | | | | |
| --- | --- | --- | --- | --- | --- | --- | --- | --- | --- | --- | --- | --- | --- | --- | --- | --- |
| Scoary | | | | | | | | | | BlastP | | | | | | |
| Gene | Number  pos  present  in (stool isolates) | Number neg present  In (stool isolates) | Number  pos not  present  in (total isolates) | Number  neg not  present  in (total isolates) | Sensitivity | Specificity | Odds  ratio | Naive  p | Bonferroni  p | Description | Max Score | Total Score | Query Cover | E value | Per. Ident | Accession |
| group_10171 | 7 | 10 | 6 | 971 | 53.84615385 | 98.98063201 | 113.2833333 | 1.70E-10 | 3.23E-06 | MULTISPECIES: DUF948 domain-containing protein [Streptococcus] | 254 | 254 | 100% | 4.00E-85 | 100.00% | WP_000569599.1 |
| group_10172 | 6 | 971 | 7 | 10 | 46.15384615 | 1.019367992 | 0.008827424 | 1.70E-10 | 3.23E-06 | MULTISPECIES: DUF948 domain-containing protein [Streptococcus] | 254 | 254 | 100% | 4.00E-85 | 100.00% | WP_000569599.1 |
| group_3927 | 5 | 1 | 8 | 980 | 38.46153846 | 99.8980632 | 612.5 | 9.58E-10 | 1.82E-05 | DUF1275 domain-containing protein [Streptococcus agalactiae] | 307 | 307 | 100% | 1.00E-105 | 100.00% | WP_025197689.1 |
| cshB | 7 | 975 | 6 | 6 | 53.84615385 | 0.611620795 | 0.007179487 | 1.16E-09 | 2.20E-05 | MULTISPECIES: DEAD/DEAH box helicase [Streptococcus] | 911 | 911 | 100% | 0 | 100.00% | WP_000008549.1 |
| group_6880 | 6 | 7 | 7 | 974 | 46.15384615 | 99.28644241 | 119.2653061 | 2.14E-09 | 4.07E-05 | DEAD/DEAH box helicase [Streptococcus agalactiae] | 912 | 912 | 100% | 0 | 100.00% | WP_000008546.1 |
| lgt | 8 | 979 | 5 | 2 | 61.53846154 | 0.203873598 | 0.003268641 | 3.33E-09 | 6.34E-05 | prolipoprotein diacylglyceryl transferase [Streptococcus agalactiae STIR-CD-09] | 517 | 517 | 100% | 0 | 100.00% | EPU03797.1 |
| group_12599 | 6 | 9 | 7 | 972 | 46.15384615 | 99.08256881 | 92.57142857 | 6.16E-09 | 0.000117217 | MULTISPECIES: Minor spike protein [Bacteria] | 662 | 662 | 100% | 0 | 100.00% | WP_000466547.1 |
| group_18006 | 6 | 10 | 7 | 971 | 46.15384615 | 98.98063201 | 83.22857143 | 9.80E-09 | 0.0001864 | DNA-binding protein [Pedobacter sp. V48] | 74.7 | 74.7 | 100% | 6.00E-17 | 100.00% | ETZ20828.1 |
| group_18007 | 6 | 10 | 7 | 971 | 46.15384615 | 98.98063201 | 83.22857143 | 9.80E-09 | 0.0001864 | C protein [Clostridioides difficile] | 98.6 | 98.6 | 100% | 5.00E-26 | 100.00% | WP_049879948.1 |
| group_2059 | 4 | 0 | 9 | 981 | 30.76923077 | 100 | inf | 1.77E-08 | 0.000336486 | leucine-rich repeat protein [Streptococcus agalactiae] | 188 | 188 | 95% | 5.00E-60 | 98.92% | WP_144061715.1 |
| group_18584 | 4 | 0 | 9 | 981 | 30.76923077 | 100 | inf | 1.77E-08 | 0.000336486 | Uncharacterised protein [Mycobacteroides abscessus subsp. abscessus] | 84.7 | 84.7 | 87% | 8.00E-18 | 52.43% | SIN58476.1 |
| group_4758 | 4 | 0 | 9 | 981 | 30.76923077 | 100 | inf | 1.77E-08 | 0.000336486 | Argininosuccinate lyase [Streptococcus agalactiae] | 243 | 243 | 100% | 3.00E-77 | 100.00% | AUP09270.1 |
| tig | 7 | 966 | 6 | 15 | 53.84615385 | 1.529051988 | 0.018115942 | 6.44E-08 | 0.001224857 | trigger factor [Streptococcus agalactiae] | 858 | 858 | 100% | 0.00E+00 | 100.00% | WP_000107756.1 |
| group_6271 | 4 | 1 | 9 | 980 | 30.76923077 | 99.8980632 | 435.5555556 | 8.78E-08 | 0.001670193 | tryptophan--tRNA ligase [Streptococcus agalactiae] | 243 | 24300% | 100% | 1.00E-79 | 100.00% | WP_025196271.1 |
| group_4829 | 4 | 1 | 9 | 980 | 30.76923077 | 99.8980632 | 435.5555556 | 8.78E-08 | 0.001670193 | CopY/TcrY family copper transport repressor [Streptococcus agalactiae] | 174 | 174 | 100% | 3.00E-54 | 100.00% | WP_060458301.1 |
| ftsH | 7 | 965 | 6 | 16 | 53.84615385 | 1.630988787 | 0.019343696 | 8.80E-08 | 0.001673833 | ATP-dependent metallopeptidase FtsH/Yme1/Tma family protein [Streptococcus agalactiae] | 1338 | 1338 | 100% | 0 | 100.00% | WP_000794531.1 |
| lepA_1 | 7 | 964 | 6 | 17 | 53.84615385 | 1.732925586 | 0.020573997 | 1.18E-07 | 0.002250677 | MULTISPECIES: translation initiation factor IF-2 [Streptococcus] | 1889 | 1889 | 100% | 0 | 99.89% | WP_000039152.1 |
| yxeM_2 | 7 | 964 | 6 | 17 | 53.84615385 | 1.732925586 | 0.020573997 | 1.18E-07 | 0.002250677 | MULTISPECIES: amino acid ABC transporter substrate-binding protein [Streptococcus] | 535 | 535 | 100% | 0 | 100.00% | WP_000473468.1 |
| ftsK | 8 | 974 | 5 | 7 | 61.53846154 | 0.713557594 | 0.011498973 | 1.21E-07 | 0.002310445 | DNA translocase FtsK [Streptococcus agalactiae] | 1645 | 1645 | 100% | 0 | 100.00% | WP_000231564.1 |
| group_6056 | 6 | 18 | 7 | 963 | 46.15384615 | 98.16513761 | 45.85714286 | 1.57E-07 | 0.002982441 | signal recognition particle protein [Streptococcus agalactiae] | 1053 | 1053 | 100% | 0 | 99.81% | WP_060457574.1 |
| group_8122 | 6 | 18 | 7 | 963 | 46.15384615 | 98.16513761 | 45.85714286 | 1.57E-07 | 0.002982441 | amino acid ABC transporter substrate-binding protein [Streptococcus agalactiae] | 535 | 535 | 100% | 0 | 100.00% | WP_000473470.1 |
| group_3923 | 5 | 922 | 8 | 59 | 38.46153846 | 6.014271152 | 0.039994577 | 2.78E-07 | 0.005288298 | DUF1275 domain-containing protein [Streptococcus agalactiae] | 466 | 466 | 100% | 4.00E-166 | 100.00% | WP_001250493.1 |
| phoU_1 | 7 | 960 | 6 | 21 | 53.84615385 | 2.140672783 | 0.025520833 | 3.38E-07 | 0.006438636 | phosphate signaling complex protein PhoU [Streptococcus agalactiae] | 441 | 441 | 100% | 1.00E-156 | 100.00% | WP_000946329.1 |
| group_9467 | 6 | 22 | 7 | 959 | 46.15384615 | 97.75739042 | 37.36363636 | 4.28E-07 | 0.008144061 | MULTISPECIES: phosphate signaling complex protein PhoU [Streptococcus] | 442 | 442 | 100% | 8.00E-157 | 100.00% | WP_000946330.1 |
| group_12598 | 5 | 10 | 8 | 971 | 38.46153846 | 98.98063201 | 60.6875 | 4.51E-07 | 0.008583529 | MULTISPECIES: scaffolding protein D [Bacteria] | 312 | 312 | 100% | 1.00E-107 | 100.00% | WP_000084700.1 |
| group_10167 | 5 | 10 | 8 | 971 | 38.46153846 | 98.98063201 | 60.6875 | 4.51E-07 | 0.008583529 | MULTISPECIES: phage capsid protein [Bacteria] | 605 | 605 | 100% | 0 | 100.00% | WP_000065548.1 |
| group_6161 | 8 | 971 | 5 | 10 | 61.53846154 | 1.019367992 | 0.016477858 | 4.51E-07 | 0.008583529 | nucleotidyltransferase [Streptococcus agalactiae] | 770 | 770 | 100% | 0 | 100.00% | WP_000218979.1 |
| gsiC | 7 | 957 | 6 | 24 | 53.84615385 | 2.44648318 | 0.029258098 | 6.66E-07 | 0.012677647 | ABC transporter permease [Streptococcus agalactiae] | 650 | 650 | 100% | 0 | 99.69% | WP_000680650.1 |
| group_3582 | 6 | 25 | 7 | 956 | 46.15384615 | 97.45158002 | 32.77714286 | 8.21E-07 | 0.015623055 | MULTISPECIES: ABC transporter permease [Streptococcus] | 649 | 649 | 100% | 0 | 99.69% | WP_000680645.1 |
| pepN | 7 | 956 | 6 | 25 | 53.84615385 | 2.54841998 | 0.030509066 | 8.21E-07 | 0.015623055 | M1 family metallopeptidase [Streptococcus agalactiae] | 1752 | 1752 | 100% | 0 | 100.00% | WP_000859371.1 |
| group_6072 | 4 | 4 | 9 | 977 | 30.76923077 | 99.5922528 | 108.5555556 | 1.20E-06 | 0.022875694 | 40S ribosomal protein S1 [Streptococcus agalactiae] | 786 | 786 | 100% | 0 | 100.00% | AKI95561.1 |
| group_4760 | 4 | 4 | 9 | 977 | 30.76923077 | 99.5922528 | 108.5555556 | 1.20E-06 | 0.022875694 | dihydroxyacetone kinase [Streptococcus agalactiae FSL S3-251] | 1120 | 1120 | 100% | 0 | 99.82% | EPV89146.1 |
| group_3898 | 5 | 13 | 8 | 968 | 38.46153846 | 98.67482161 | 46.53846154 | 1.26E-06 | 0.023994525 | MULTISPECIES: NADH oxidase [Streptococcus] | 939 | 939 | 100% | 0 | 100.00% | WP_000036813.1 |
| carB_1 | 5 | 13 | 8 | 968 | 38.46153846 | 98.67482161 | 46.53846154 | 1.26E-06 | 0.023994525 | carbamoyl phosphate synthase large subunit [Streptococcus agalactiae] | 823 | 823 | 100% | 0 | 100.00% | RDY91281.1 |
| group_2749 | 8 | 968 | 5 | 13 | 61.53846154 | 1.325178389 | 0.021487603 | 1.26E-06 | 0.023994525 | insulinase family protein [Streptococcus agalactiae] | 841 | 841 | 100% | 0 | 100.00% | WP_000706191.1 |
| group_5238 | 6 | 28 | 7 | 953 | 46.15384615 | 97.14576962 | 29.17346939 | 1.47E-06 | 0.028010324 | MULTISPECIES: MurR/RpiR family transcriptional regulator [Streptococcus] | 554 | 554 | 100% | 0 | 100.00% | WP_000158188.1 |
| group_10272 | 5 | 14 | 8 | 967 | 38.46153846 | 98.57288481 | 43.16964286 | 1.70E-06 | 0.032342565 | MULTISPECIES: putative DNA-binding protein [Streptococcus] | 218 | 218 | 100% | 9.00E-72 | 100.00% | WP_000402075.1 |
| group_10271 | 8 | 967 | 5 | 14 | 61.53846154 | 1.427115189 | 0.023164426 | 1.70E-06 | 0.032342565 | MULTISPECIES: putative DNA-binding protein [Streptococcus] | 218 | 218 | 100% | 9.00E-72 | 100.00% | WP_000402075.1 |
| group_2390 | 3 | 0 | 10 | 981 | 23.07692308 | 100 | inf | 1.75E-06 | 0.033345735 | Membrane protein [Streptococcus agalactiae] | 577 | 577 | 100% | 0 | 100.00% | AKI58410.1 |
| group_736 | 3 | 0 | 10 | 981 | 23.07692308 | 100 | inf | 1.75E-06 | 0.033345735 | phosphoribosylformylglycinamidine synthase [Streptococcus agalactiae] | 1574 | 1574 | 100% | 0 | 100.00% | CNC33084.1 |
| group_3769 | 3 | 0 | 10 | 981 | 23.07692308 | 100 | inf | 1.75E-06 | 0.033345735 | hypothetical protein TH70_0672 [Streptococcus agalactiae] | 208 | 208 | 100% | 2.00E-65 | 100.00% | ODG95866.1 |
| group_5676 | 3 | 0 | 10 | 981 | 23.07692308 | 100 | inf | 1.75E-06 | 0.033345735 | conserved hypothetical protein [Streptococcus agalactiae 515] | 168 | 168 | 100% | 7.00E-53 | 100.00% | EAO70900.1 |
| group_8728 | 3 | 0 | 10 | 981 | 23.07692308 | 100 | inf | 1.75E-06 | 0.033345735 | metal ABC transporter substrate-binding protein [Streptococcus agalactiae] | 625 | 625 | 100% | 0 | 99.68% | WP_079859693.1 |
| group_2028 | 3 | 0 | 10 | 981 | 23.07692308 | 100 | inf | 1.75E-06 | 0.033345735 | bifunctional phosphoribosylaminoimidazolecarboxamide formyltransferase/IMP cyclohydrolase [Streptococcus agalactiae] | 438 | 438 | 99% | 8.00E-153 | 100.00% | WP_025197374.1 |
| group_5545 | 3 | 0 | 10 | 981 | 23.07692308 | 100 | inf | 1.75E-06 | 0.033345735 | ATP-dependent Clp protease ATP-binding subunit [Streptococcus agalactiae] | 879 | 879 | 100% | 0 | 100.00% | WP_025196792.1 |
| group_2751 | 3 | 0 | 10 | 981 | 23.07692308 | 100 | inf | 1.75E-06 | 0.033345735 | insulinase family protein [Streptococcus agalactiae] | 753 | 753 | 100% | 0 | 99.73% | WP_115341608.1 |
| group_3197 | 3 | 0 | 10 | 981 | 23.07692308 | 100 | inf | 1.75E-06 | 0.033345735 | argininosuccinate synthase [Streptococcus agalactiae] | 816 | 816 | 100% | 0 | 100.00% | WP_000514045.1 |
| group_9496 | 3 | 0 | 10 | 981 | 23.07692308 | 100 | inf | 1.75E-06 | 0.033345735 | deoxyribose-phosphate aldolase [Streptococcus agalactiae] | 105 | 105 | 100% | 3.00E-27 | 100.00% | WP_101791143.1 |
| group_7947 | 6 | 29 | 7 | 952 | 46.15384615 | 97.04383282 | 28.13793103 | 1.77E-06 | 0.033595842 | MULTISPECIES: arginine repressor [Streptococcus] | 333 | 333 | 100% | 9.00E-116 | 100.00% | WP_001034427.1 |
| group_8311 | 4 | 5 | 9 | 976 | 30.76923077 | 99.490316 | 86.75555556 | 2.15E-06 | 0.040876144 | Inhibitor of apoptosis-promoting Bax1 [Streptococcus agalactiae] | 134 | 134 | 100% | 2.00E-39 | 100.00% | CNK78606.1 |
| group_9821 | 9 | 113 | 4 | 868 | 69.23076923 | 88.48114169 | 17.28318584 | 2.24E-06 | 0.042632717 | site-specific integrase [Streptococcus agalactiae] | 353 | 353 | 100% | 7.00E-123 | 100.00% | WP_025193995.1 |
| proB | 8 | 966 | 5 | 15 | 61.53846154 | 1.529051988 | 0.02484472 | 2.25E-06 | 0.042829987 | glutamate 5-kinase [Streptococcus agalactiae] | 539 | 539 | 100% | 0 | 99.63% | WP_001874749.1 |
| group_3606 | 7 | 950 | 6 | 31 | 53.84615385 | 3.160040775 | 0.038070175 | 2.50E-06 | 0.047522061 | PTS cellobiose transporter subunit IIC [Streptococcus agalactiae] | 920 | 920 | 100% | 0 | 100.00% | WP_000085851.1 |
